# Recurrent Loss of APOBEC3H Activity during Primate Evolution

**DOI:** 10.1101/311878

**Authors:** Erin I. Garcia, Michael Emerman

## Abstract

Genes in the *APOBEC3* family encode cytidine deaminases that provide a barrier against viral infection and retrotransposition. Of all *APOBEC3* genes in humans, *APOBEC3H (A3H)* is the most polymorphic: some haplotypes encode stable and active A3H proteins, while others are unstable and inactive. Such variation in human A3H affects interactions with the lentiviral antagonist Vif, which counteracts A3H via proteasomal degradation. In order to broaden our understanding of A3H-Vif interactions as well as its evolution in Old World monkeys, we characterized A3H variation within four African green monkey (AGM) subspecies. We found that A3H is highly polymorphic in AGMs and has lost antiviral activity in multiple Old World monkeys. This loss of function was partially related to protein expression levels but was also influenced by amino acid mutations in the N-terminus. Moreover, we demonstrate that the evolution of A3H in the primate lineages leading to AGMs was not driven by Vif. Our work suggests that activity of A3H is evolutionarily dynamic and may have a negative effect on host fitness, resulting in its recurrent loss in primates.

**Importance:** Adaptation of viruses to their hosts is critical for transmission of viruses between different species. Previous studies had identified changes in a protein from the APOBEC3 family that influenced species-specificity of simian immunodeficiency viruses (SIVs) in African green monkeys. We studied the evolution of a related protein in the same system, APOBEC3H, which has experienced a loss of function in humans. This evolutionary approach revealed that recurrent loss of APOBEC3H activity has taken place during primate evolution suggesting that APOBEC3H places a fitness cost on hosts. The variability of APOBEC3H activity between different primates highlights the differential selective pressures on the *APOBEC3* gene family.

## Introduction

The seven members of the *APOBEC3 (A3)* gene family in primates encode cytidine deaminases involved in innate immune defense against retroviruses and retroelements (1–3). Four A3 enzymes are known to potently restrict the replication of lentiviruses like simian immunodeficiency virus (SIV) or human immunodeficiency virus (HIV): A3D, A3F, A3G, and A3H (4). A3 proteins are packaged into budding virions and cause G-to-A hypermutation of viral DNA, although deamination-independent modes of restriction have also been characterized (5). Hypermutation of viral DNA results in detrimental mutations that render the virus inactive and thus protects new cells from infection. However, lentiviruses have evolved a mechanism to evade A3 restriction by encoding a viral antagonist, Vif, which binds and targets A3 proteins for proteasomal degradation via a cellular E3 ubiquitin ligase complex (6). Given that A3-Vif interactions often act in a species-specific manner, adaptation of Vif to host A3 proteins is important for successful adaptation of lentiviruses to their hosts (7–11).

Various A3 proteins differ in their ability to restrict viral infection. For instance, A3G is the most potent A3-mediated inhibitor of HIV-1, while A3A and A3B have limited antiviral potential (4, 12). In humans, the A3H protein is especially remarkable because multiple polymorphisms drastically impact its antiviral activity (13–16). Two independent mutations have occurred in human evolution that destabilized A3H (13); haplotypes that encode an R105G mutation or a deletion of amino acid 15 make unstable proteins (haplotypes I, III, IV, VI) that have lost antiviral activity, while those without these changes make stable proteins (haplotypes II, V, VII) that potently restrict HIV-1 (13-15, 17). Stability and antiviral activity of human A3H has been further linked to subcellular localization: unstable/inactive proteins are more nuclear, while stable/active proteins remain cytoplasmic (18). A3H haplotype I has been associated with breast and lung cancer (19), as has another nuclear A3, A3B (20), suggesting that in some cases A3 activity may be detrimental for the host. While such events have occurred in humans, examples of gains or losses of A3 activity over evolutionary time in other primates have been less explored.

The *A3H* genetic polymorphisms present in human populations also impact the interactions between human A3H and HIV-1 Vif (17, 21–23). Stable A3H proteins are only partially susceptible to degradation by Vif from the LAI isolate of HIV-1 and not at all by HIV-1 NL4-3 Vif (21, 23). However, studies using human cohorts encoding different haplotypes of *A3H* have shown that Vif proteins from primary virus strains isolated from patients with *A3H* haplotype II are able to antagonize stable proteins, while those from patients with unstable haplotypes cannot. This suggests that there is selection *in vivo* for Vif strains that counteract the stably expressed forms of A3H in infected people (22, 24, 25). Furthermore, the cross-species transmissions that led to adaptation of SIV from monkeys to chimpanzees to humans, giving rise to HIV-1, involved adaptation of Vif to antagonize the A3 proteins found in each host (7, 8, 10, 11) including adaptation of Vif from SIVcpz to antagonize human A3H (11).

The study of the evolution of lentiviral-host interactions within Old World monkeys has provided insights into the longer-term dynamics of the evolutionary arms race between host antiviral proteins and their lentiviral targets (7, 9, 26). African green monkeys (AGMs), in particular, provide a unique opportunity to assess the evolutionary forces governing interactions between lentiviruses and their hosts since the genus *Chlorocebus* encompasses four geographically distinct subspecies (vervets, sabaeus, tantalus, and grivets) each of which are infected with species-specific subtypes of SIVagm (27–29). These SIV subtypes are named by the subspecies they infect: SIVagm.ver, SIVagm.sab, SIVagm.tan, and SIVagm.gri (29). Furthermore, this system is particularly powerful because although these subspecies are closely genetically related, enough divergence exists that has allowed species-specific lentiviral infections to occur.

Previous studies have demonstrated that many genes involved in antiviral immunity are polymorphic in AGMs and some changes at the protein level are critical for interactions with viral antagonists (8, 30, 31). A3G in particular was discovered to encode species-specific polymorphisms (8). Amino acid changes in the Vif binding domain of AGM A3G (sites 128 and 130) confer protection against SIVagm strains from other subspecies. For example, K128E, found in grivet monkeys, is resistant to all Vifs except SIVagm.gri, while D130H, found in sabaeus monkeys, is resistant to Vifs from SIVagm.ver and SIVagm.tan (8). This demonstrates that Vif continues to drive the evolution of A3G and contributes to the species-specificity of lentiviruses in AGM populations. It is unknown whether A3H has a similar role in these primates.

In this study, we asked whether we could identify a host-virus “arms-race” between A3H in AGMs and Vif proteins encoded by the SIVs that infect these species. Surprisingly, we found that although A3H is highly polymorphic in AGMs, the antiviral activity of A3H has been largely lost in AGMs. The reduced antiviral activity of AGM A3H is in part caused by lower protein expression levels, although amino acid changes also lower antiviral activity independent of protein levels. By reconstructing ancestral A3H proteins spanning evolution in AGMs and in other Old World monkeys, we find that there has been recurrent loss of A3H activity in some, but not all, Old World monkeys. While higher expression levels generally increase viral inhibition, we also identified amino acids that affect A3H restriction without increasing protein expression, which map to regions implicated in RNA binding (32–34). Thus, our data support a model where A3H antiviral activity has been repeatedly lost throughout evolution. This argues that there is a longer scale dynamic between the cost and benefit for A3H function in primates that is not necessarily driven by interactions with its antagonist.

## Results

### A3H is highly polymorphic in AGM subspecies, but all tested alleles have low antiviral activity

Polymorphisms in human *A3H* are known to affect its antiviral activity as well as its interactions with Vif (13, 15, 17, 22, 23). Similarly, polymorphisms in *A3G* in AGMs impact interactions with Vif, suggesting an ongoing and ancient genetic conflict between A3G and SIV Vif in AGMs (8). These observations prompted us to explore the genetic landscape of A3H in African green monkeys and other Old World monkeys to determine if A3H evolution has been driven by Vif over a broad evolutionary scale. First, we sequenced *A3H* in 50 AGM samples collected from all four subspecies infected with a species-specific SIV: vervet, sabaeus, tantalus, and grivet monkeys. The mitochondrial DNA of these animals had also been previously sequenced and was confirmed to cluster by AGM subspecies (8). Sequence analysis of 80 independent *A3H* genes revealed 34 single-nucleotide polymorphisms (SNPs), 6 of which are synonymous changes at a frequency higher than 5%. 28 SNPs are nonsynonymous; 23 are found in more than one individual (Table 1). Most polymorphisms are represented in all four subspecies (Figure 1A and B), except for low frequency variants only found in one AGM and two nonsynonymous mutations found in two vervet sequences at amino acid residues 113 and 116. Phylogenetic analysis showed that *A3H* sequences from all AGM subspecies are paraphyletic (Figure 1A), similar to a previous study examining *A3G* in these species (8).

**Figure 1.**
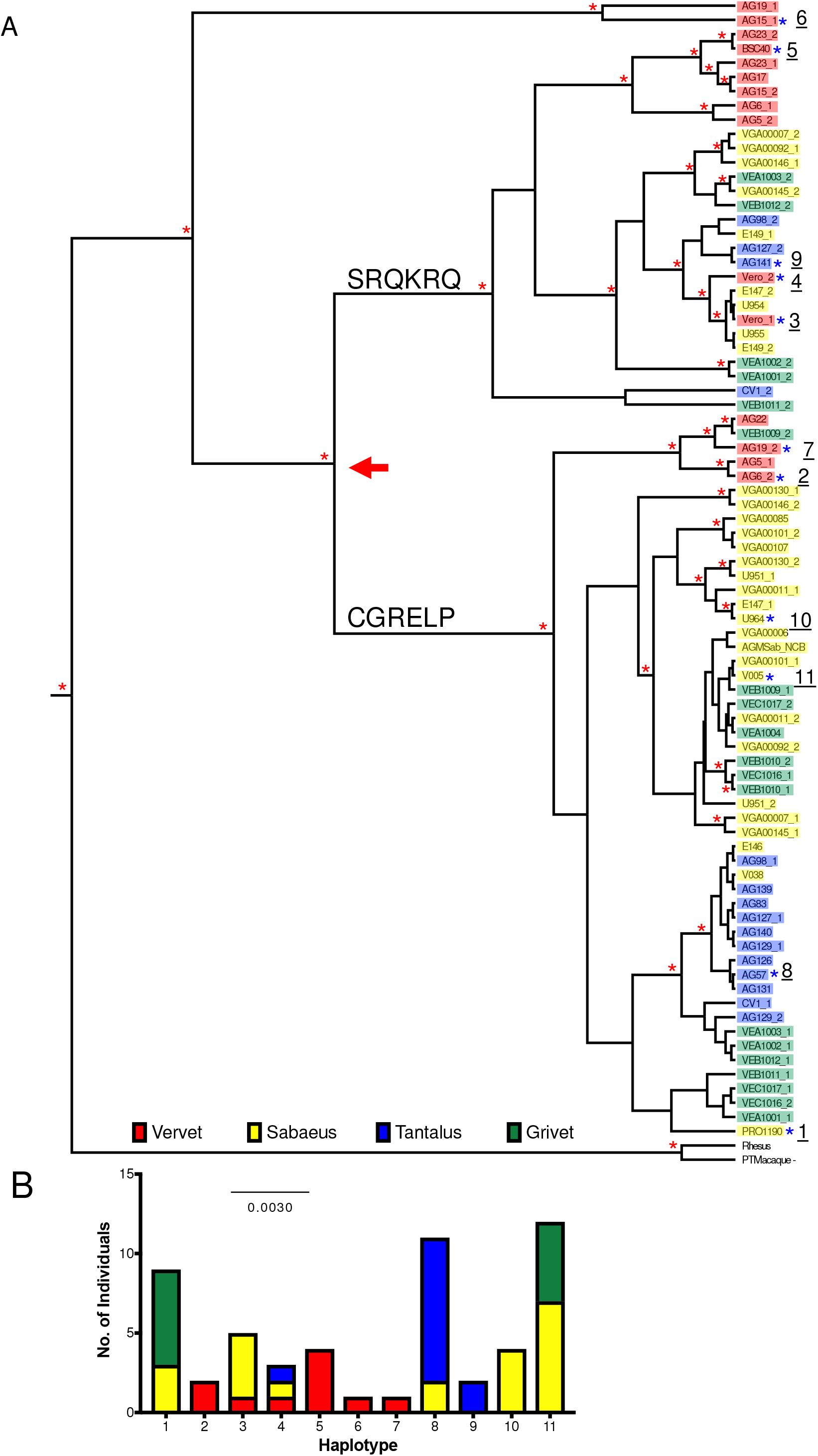
Sequence and Phylogenetic Analysis of *A3H* in African green monkeys. **A**. The evolutionary relationship between 80 full-length AGM *A3H* genes was inferred by Bayesian MCMC phylogenetic reconstruction. Red asterisks (*) show nodes that have a posterior probability > 0.5. Colored boxes demonstrate the subspecies of origin (red = Vervet, yellow = Sabaeus, blue = Tantalus, and green = Grivet). Blue asterisks (*) denote cloned haplotypes, numbered 1 – 11. **B**. Number of AGM individuals encoding haplotypes 1 – 11, color coded by subspecies similar to *A3H* phylogeny.

**Table 1.** Polymorphisms identified in AGM *A3H* sequences.

| Site | Observed amino acids | Predicted location in structure<sup>s</sup> |
| --- | --- | --- |
| 17 | R/H | Loop 1 |
| 18 | Y/R/H/L* |  |
| 20 | S/N |  |
| 25 | P/R |  |
| 41 | T/M* | Loop 2 |
| 44 | R/K* | β-sheet |
| 46 | H/Q |  |
| 53 | H/D | Loop 3 |
| 73 | C/S* | Loop 4 |
| 75 | R/Q | β-sheet |
| 113 | Y/C | Loop 7 |
| 116 | R/H |  |
| 117 | R/P* |  |
| 127 | C/S | α4 |
| 128 | G/R |  |
| 130 | R/Q | Loop 8 |
| 134 | E/K | β-sheet |
| 138 | L/R | α5 |
| 139 | P/Q |  |
| 152 | K/E | Loop 10 |
| 164 | D/E | α6 |
| 171 | Q/R |  |
| 181 | K/E |  |
| 195 | N/S | Not resolved in structure |
| 204 | S/A |  |

\* Amino acid residues only identified in one animal
<sup>a</sup>Location in AGM A3H modeled onto a previously described pig-tailed macaque A3H (Bohn et al, PDB 5W3V)

Nonsynonymous SNPs are spread throughout the protein (Table 1), although one group is clustered between amino acids 127 – 139. The changes are tightly linked and result in divergence of the phylogenetic tree between two clades (Figure 1A, red arrow). Haplotypes with a CGRELP motif compose one clade on the tree, while the other has an SRQKRQ motif. Residues 127 and 128 are located on the α4 helix, which has been implicated in A3H-Vif binding (35, 36). Furthermore, the regions of *A3H* with the greatest number of nonsynonymous mutations are in the predicted loops 1 and 7, and the α6 helix (Table 1), which were recently shown by structural studies to be involved in an interaction between A3H and a co-crystalized RNA (32–34). These results indicate that, similar to A3G, *SAMHD1,* and other antiviral genes in AGMs (7, 30, 31), there is extensive polymorphism in AGM *A3H.*

The presence of numerous polymorphisms suggests there may be functional consequences for either antiviral activity or Vif antagonism. To determine whether or not nonsynonymous polymorphisms in *A3H* impact antiviral activity, we tested the ability of A3H protein variants to restrict lentiviruses. We cloned haplotypes, numbered 1 – 11, which are representative of each clade in the phylogenetic tree into a mammalian expression vector for functional analysis (Figure 1B). These haplotypes also represent all four subspecies of AGMs. Out of the 28 total nonsynonymous SNPs found in AGMs (Table 1), 11 unique protein sequences were tested and 20 polymorphic sites were characterized (Table 2).

**Table 2.**
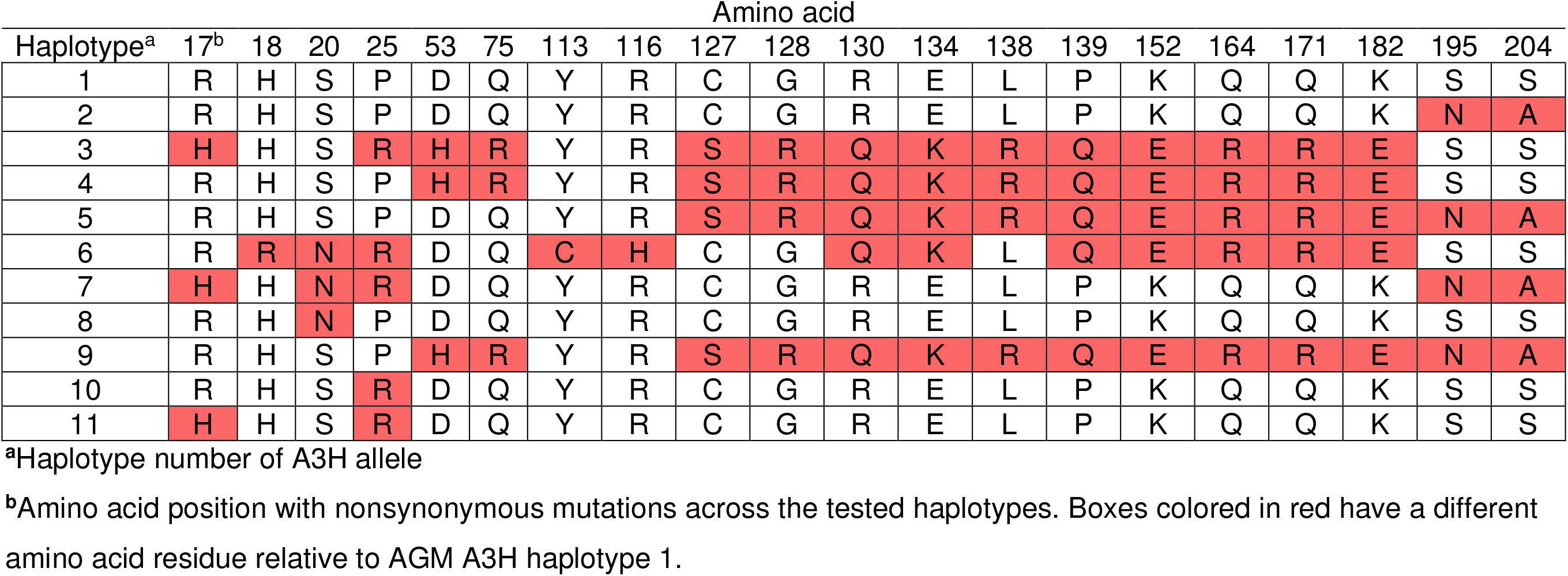
Polymorphic sites in tested A3H haplotypes.

A3H expressing plasmids were co-transfected into HEK293T cells for singleround infectivity experiments with *HIVΔenvΔvif* or SIVagm.TAN *ΔenvΔvif* proviruses and a VSV-G expression plasmid for pseudotyping to measure antiviral activity. Viral supernatants were collected, normalized for RT activity, and used to infect SupT1 cells. We found that none of the AGM A3H variants restricted lentiviruses as potently as the A3H from rhesus macaque (Figure 2A), an Old World monkey that has been previously characterized for its A3H activity (4, 13, 14). Rhesus A3H restricted viral infection of HIVΔ*vif* by approximately 21-fold, while AGM A3H variants restrict viral infection by no more than 3-fold, and some not at all (Figure 2A). Although activity against HIV-1 *Δvif* correlates with activity against other lentiviruses, we also validated this result using SIVagm.TANΔ*vif*, a strain originally isolated from tantalus monkeys. We found that the AGM A3H haplotypes were also poorly restrictive against SIVagm.TANΔ*vif* compared to the activity of rhesus macaque A3H (Figure 2A). This indicates A3H variants encoded by AGMs have poor antiviral activity against at least two separate lentiviruses and is not due to species-specificity (Figure 2B).

**Figure 2.**
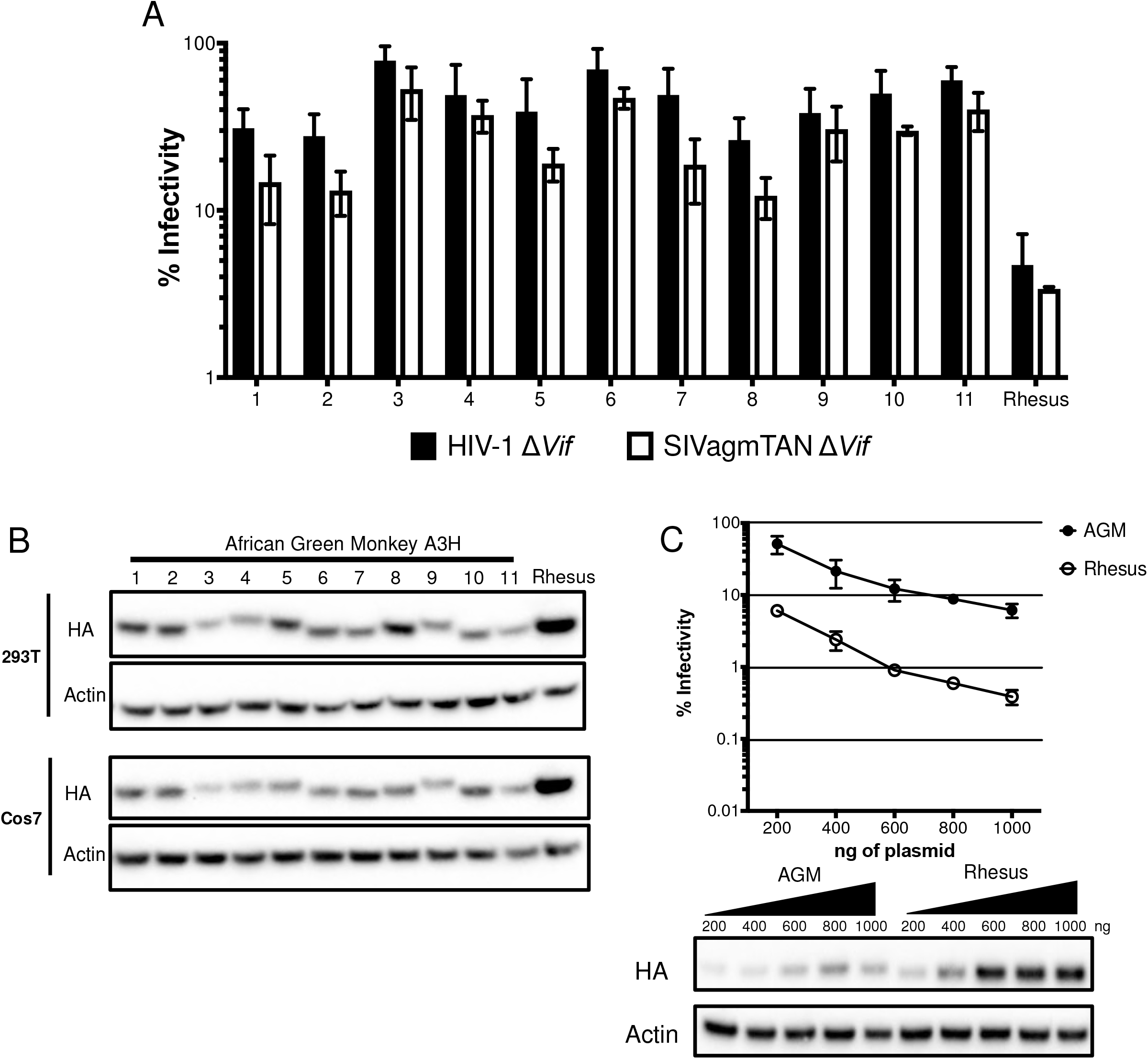
Antiviral activity of A3H is lower in AGMs than in another Old World monkey. **A**. Single-cycle infectivity assays were performed in the presence or absence of A3 proteins against HIVΔvif (black) and SIVagmΔvif (white). Rhesus macaque was included as a positive control. Relative infection was normalized to viral infectivity in the absence of A3 proteins. Averages of three replicates, each with triplicate infections ( ± SEM) are shown. **B**. Western blot analysis of HA-tagged AGM A3H protein expression in human (HEK293T) and AGM (Cos7) cell lines. The different size bands for different AGM A3H haplotypes is reproducible. β-Actin is shown as a loading control. **C**. Top: Single-cycle infectivity assay of *HIVΔvif* in the presence of increasing amounts of A3H-expressing plasmids. AGM A3H haplotype 1 (black circles) and rhesus macaque A3H (open circles) are compared. Relative infection was normalized to viral infectivity in the absence of A3 proteins. Averages of three replicates, each with triplicate infections ( ± SEM) are shown. Bottom: Western blot analysis of protein expression levels with the same amounts of plasmid in the top panel. β-Actin is shown as a loading control.

The differences in restriction could be explained by changes in expression since we found that no variant of AGM A3H is as strongly expressed as rhesus macaque A3H in HEK293T cells (Figure 2B, top). This lower expression of AGM A3H proteins relative to rhesus macaque A3H was not due to the species of origin of the cells used for transfection since when we transfected AGM-derived Cos7 cells, the protein expression levels of AGM A3H were similarly poor relative to the expression of the rhesus A3H (Figure 2B, bottom). These data indicate that lower protein expression levels are correlated with less potent antiviral activity, similar to unstable haplotypes of human A3H.

To ask whether lower antiviral activity of AGM A3H proteins was due to low expression levels alone, we selected one of the most potent AGM A3H proteins, haplotype 1 (Figure 2A), and increased the amount of transfected plasmid from 200ng to 1000ng in parallel to rhesus macaque A3H. At the highest plasmid concentration, haplotype 1 was able to restrict HIV*Δvif* 16-fold, while rhesus A3H restricted HIVΔ*vif* 19-fold at the lowest concentration used (Figure 2C, top). This corresponds to approximately equal levels of protein expression at these amounts of plasmid transfected (Figure 2C, bottom; compare band for AGM A3H at the highest level of plasmid transfected with the level of Rhesus A3H at the lowest level of plasmid transfected). This demonstrates that AGM A3H has not inherently lost its function, since as the protein concentration increases, the antiviral activity correspondingly becomes more potent.

In order to more fully explore the relationship between expression levels and antiviral activity of AGM A3H, we codon-optimized both AGM *A3H* haplotype 1 and rhesus macaque *A3H* sequences to remove rare codons that might negatively affect protein translation efficiency. Based on codon-usage statistics in primates (see Methods) 100 out of 211 codons were replaced to more frequent codons in the codon-optimized AGM haplotype 1 *A3H* and 106 out of 211 were replaced in the codon-optimized rhesus macaque *A3H.* We found that codon-optimization of both AGM A3H and of rhesus macaque A3H increased their expression levels relative to the native codons in each gene (Figure 3A—compare WT to CO (codon-optimized)). Moreover, after codon-optimization, AGM A3H haplotype 1 and rhesus macaque A3H are expressed to similar levels (Figure 3A and Figure 3B, bottom). However, while codon optimization increased the antiviral activity of both AGM and rhesus macaque A3H over that of wild-type (compare Figure 3B to 2D), codon-optimized rhesus macaque A3H still restricts viral infection 10-fold better than codon-optimized AGM A3H (Figure 3B, top). This shows that while increasing the expression level of the protein is sufficient to improve its antiviral activity, other factors also influence protein function.

**Figure 3.**
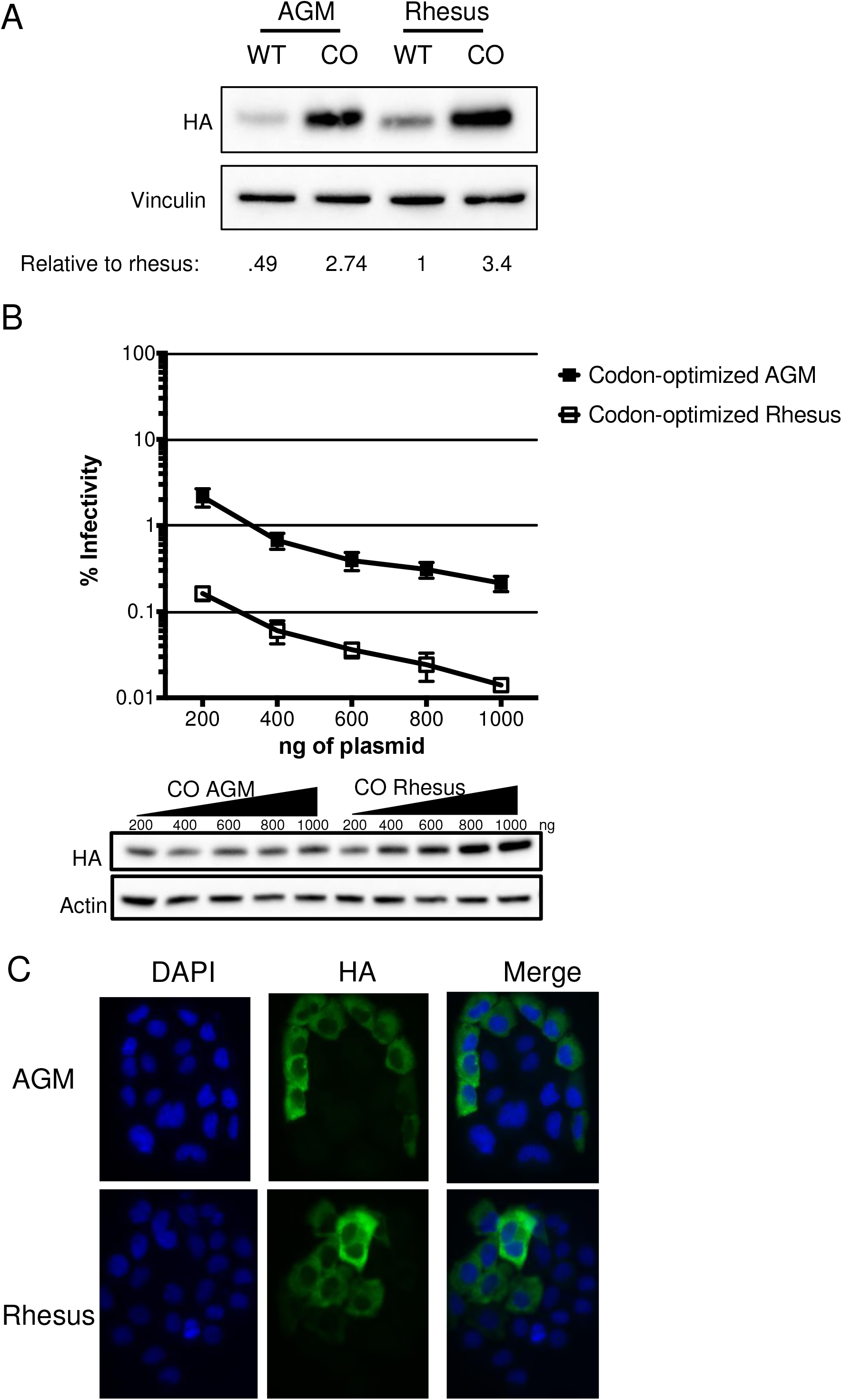
Codon-optimization increases protein expression and antiviral activity. **A**. Western blot analysis for the expression of AGM A3H haplotype 1, codon-optimized haplotype 1 A3H, rhesus macaque A3H, and codon-optimized rhesus macaque A3H. Vinculin was used as a protein loading control. Quantification was done relative to rhesus macaque A3H (normalized to 1). **B**. Top: Single-cycle infectivity assay of *HIVΔvif* in the presence of increasing amounts of A3H plasmid comparing codon-optimized AGM haplotype 1 A3H (black squares) and codon-optimized rhesus macaque A3H (open squares). Relative infection was normalized to viral infectivity in the absence of A3 proteins. Averages of three replicates, each with triplicate infections (± SEM) are shown. Bottom: Western blot analysis of protein expression level with amounts of plasmid added in panel B. β-Actin is shown as a loading control. **C**. Subcellular localization of rhesus macaque and AGM A3H haplotype 1 in HeLa cells. A3H proteins were detected with an anti-HA antibody (green) and DAPI staining was used to detect the nucleus (blue). Image is representative of n = 90 total images over 3 replicates.

Studies in humans have shown that inactive A3H proteins also localize to the nucleus (13, 18). We therefore asked whether AGM A3H proteins were expressed to lower levels with low antiviral activity due to a change in localization. WT AGM A3H haplotype 1 and WT rhesus macaque A3H were transfected into HeLa cells and visualized using immunofluorescent microscopy. However, both AGM and rhesus macaque A3H were mainly present in the cytoplasm (Figure 3C), demonstrating that a drastic change in localization was not responsible for the lack of potent antiviral activity of AGM A3H.

Our results suggest that the extensive diversity observed in AGM A3H results in lower antiviral activity linked to both protein levels and to other functional differences due to amino acid divergence between species.

### Reconstruction of ancestral A3H proteins demonstrates the loss of activity in more recent evolution of AGMs and other primates

We previously explored the evolutionary dynamic of hominoid A3H by reconstructing the ancestor of human/chimpanzee A3H and found that the predicted A3H protein at the human/chimpanzee ancestor had higher antiviral activity than either the extant chimpanzee or human proteins (13). This suggests that there has been a loss of some activity in both lineages over their evolution. Due to the finding that all tested AGM A3H proteins are poorly active relative to the rhesus macaque A3H, we wanted to reconstruct the ancestral history of A3H leading to the modern AGM lineage. Thus, in order to gain statistical power in the ancestral sequence predictions at each node, we determined the *A3H* sequence from a broader panel of Old World monkeys in sister clades including De Brazza’s monkey, Allen’s Swamp monkey, Wolf’s guenon, mustached guenon, talapoin, and patas monkey (Figure 4A).

**Figure 4.**
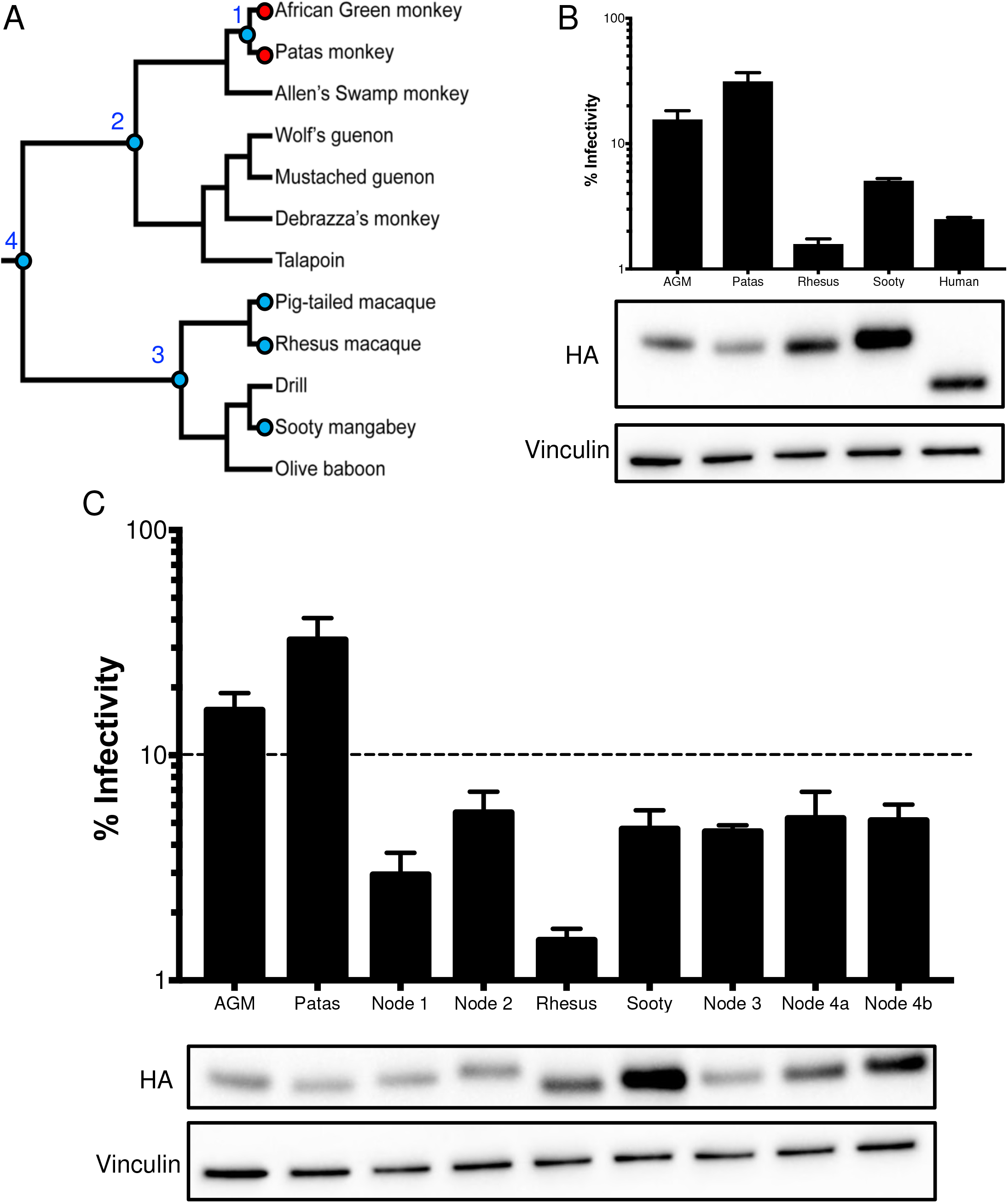
AGM ancestors encode potent antiviral proteins. **A**. A phylogeny, depicted as a cladogram, based on the accepted species tree of all sequenced Old World primates included in the study (27). Blue circles denote active antiviral proteins; red circles denote inactive antiviral proteins. Ancestral nodes are labeled with numbers (1 – 4). **B**. Top: Single-cycle infectivity assay for HIVΔvif against extant primate A3H proteins. Relative infection was normalized to viral infectivity in the absence of A3 proteins. Averages of three replicates, each with triplicate infections (± SEM) are shown. Bottom: Western blot analysis of the protein expression level. Vinculin is used as a loading control. **C**. Top: Single-cycle infectivity assay for *HIVΔvif* against ancestral A3H proteins and their extant descendants. Relative infection was normalized to viral infectivity in the absence of A3 proteins. Averages of three replicates, each with triplicate infections ( ± SEM) are shown. Dotted line at 10% is an arbitrary reference point. Bottom: Western blot analysis of the protein expression level. Vinculin is used as a loading control.

We tested the A3H activity of the closest sister species to the AGMs, patas monkeys, and the A3H activity of a sister species to the rhesus, the sooty mangabey (Figure 4A). Upon transfection into HEK293T cells, the protein expression level of patas monkey A3H was lower in comparison to rhesus macaque A3H, similar to AGM A3H (Figure 4B, bottom). Patas monkey A3H correspondingly had low antiviral activity when tested against HIVΔ*vif*; that is, while AGM and rhesus macaque A3H restricted viral infection 6-fold and 63-fold, respectively, patas monkey A3H restricted *HIVΔvif* infection only 3-fold (Figure 4B, top). However, active A3H proteins from sooty mangabey and the human A3H haplotype II restrict viral infection 20-fold and 40-fold, respectively. These data show that the antiviral activity is low in a species closely related to AGMs, but a relative of the rhesus macaque and humans encode more active A3H proteins. This finding further suggests that changes in A3H activity may have deeper evolutionary origins in primate evolution.

In order to determine whether A3H antiviral activity was gained in the rhesus macaque/sooty mangabey lineage, or was lost in the AGM/patas monkey lineage, we reconstructed the *A3H* ancestors at various nodes in the Old World monkey phylogeny (Figure 4A). These included the common ancestor of AGMs and patas monkeys (node 1), AGMs, patas monkeys, and its sister clade (node 2), as well as the common ancestor of rhesus macaque, pig-tailed macaque, and sooty mangabey (node 3) and common ancestor of both groups (node 4). Each ancestor was constructed using maximum likelihood with FastML (37) based on the primate species phylogeny.

Although the majority of codons were reconstructed with a probability >99%, site 207 was ambiguous in the common ancestor of all tested Old World monkeys and two codons were possible – encoding either an isoleucine (node 4a) or a threonine (node 4b) at position 207. In this case, both possible ancestors were generated using point-mutagenesis.

We then tested the predicted A3H protein at the reconstructed ancestral nodes of the Old World monkey phylogenetic tree for antiviral activity and protein expression level. All ancestors inhibited viral infection between 17- and 33-fold (Figure 4C). Thus, this result suggests that activity was lost within the AGM/patas monkey clade rather than specifically gained in the rhesus macaque/sooty mangabey clade. Moreover, the node 1 ancestor representing the common ancestor of AGM and patas monkeys restricted viral infection 33-fold, which was more potent than its descendants, AGMs (6-fold) and patas monkeys (3-fold), despite having similar protein expression levels (Figure 4C). Therefore, loss of activity in A3H in AGM and patas monkeys that occurred after the common ancestor at node 1, which diverged at least 4 million years ago (27), included mutations that both decreased protein expression levels and led to the losses of antiviral activity that are independent of protein expression.

### Multiple amino acid mutations are responsible for loss of A3H antiviral activity

The ancestral A3H protein at node 1 representing the common ancestor of AGMs and patas monkeys (Figure 4A) has stronger antiviral activity than its extant descendants (Figure 4C). Therefore, we wanted to trace the amino acid mutations that resulted in the subsequent loss along the branches leading from node 1 to AGMs and patas monkeys. The sequence alignment of the node 1 ancestor with AGM haplotype 1 and patas monkey A3H reveals 4 (sites 18, 20, 48, and 171) and 3 (sites 17, 25, and 51) amino acid differences respectively (Figure 5A). One site in AGMs, S20N, created a protein sequence identical to AGM A3H haplotype 8. We introduced the other residues found in the node 1 ancestor into AGM and patas monkey A3H backgrounds to test the expression level and antiviral activity of each mutant.

**Figure 5.**
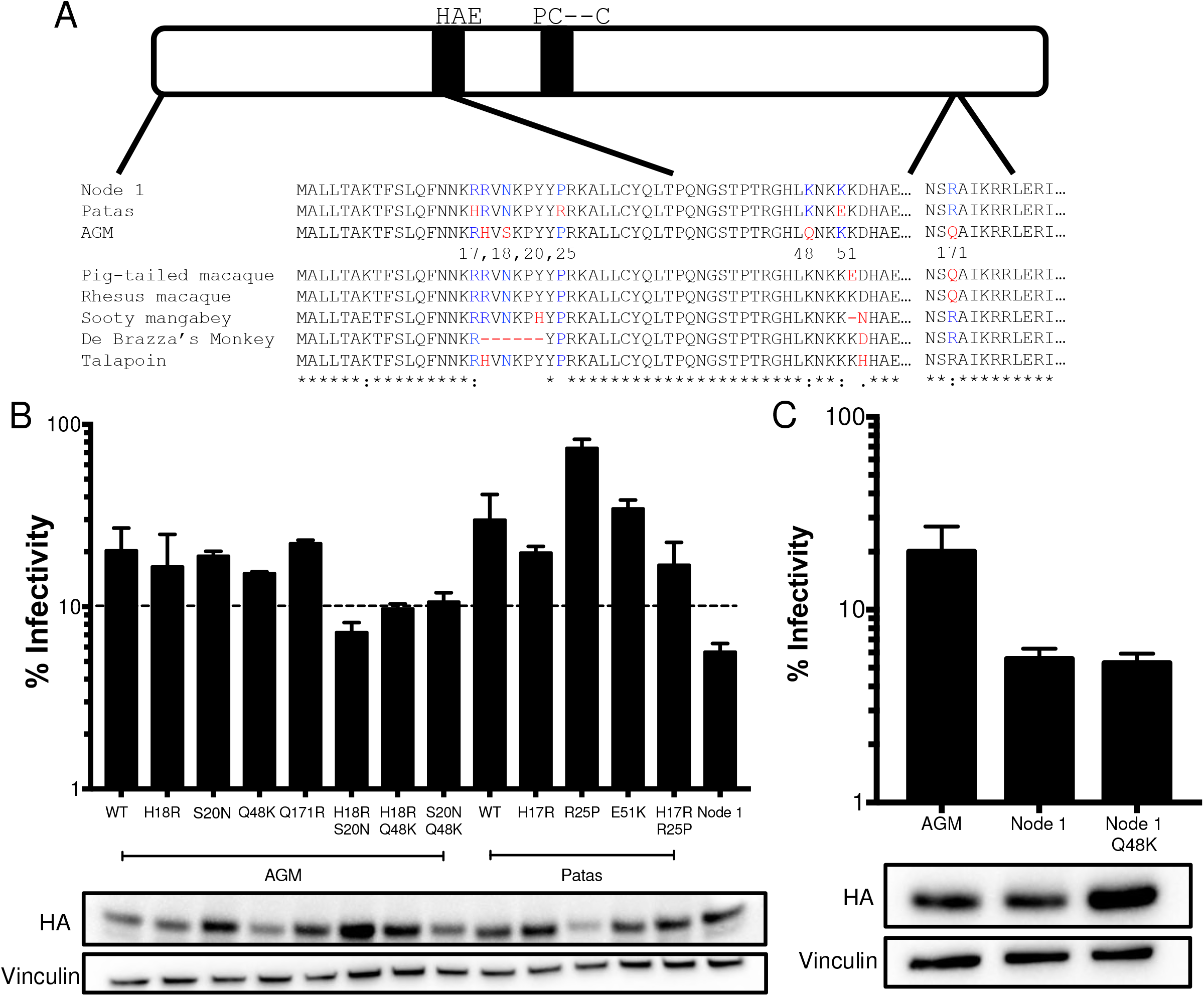
Multiple amino acid mutations required for an increase in antiviral activity. **A**. Schematic of the A3H protein. Black bars outline the A3H catalytic site. Numbered amino acid residues that are different between AGM A3H, patas A3H, and the node 1 ancestor are outlined in red on the protein sequence alignment, ancestral residues are colored blue. **B**. Top: Single-cycle infectivity assay for *HIVΔvif* against extant mutants. Relative infection was normalized to viral infectivity in the absence of A3 proteins. Averages of two replicates, each with triplicate infections ( ± SEM) are shown. Bottom: Western blot analysis of protein expression levels of HA-tagged extant mutants made in the AGM and patas A3H backgrounds. Vinculin is used as a loading control. **C**. Top: Single-cycle infectivity assay for *HIVΔvif* against AGM haplotype 1, node 1 ancestor, and node 1 mutant A3H. Relative infection was normalized to viral infectivity in the absence of A3 proteins. Averages of two replicates, each with triplicate infections (± SEM) are shown. Dotted line at 10% is an arbitrary reference point. Bottom: Western blot analysis of protein expression levels of HA-tagged proteins made in the AGM and patas A3H backgrounds. Vinculin is used as a loading control. **D**. A partial species phylogeny, shown as a cladogram, accompanied by amino acid residues across Old World monkeys at sites found to be important for antiviral activity.

Notably, no single amino acid mutation in AGM A3H increased the antiviral activity compared to wild-type AGM A3H and all single mutants inhibited viral infection around 5-fold relative to the no A3 control (Figure 5B). In contrast, the double mutants H18R/S20N, H18R/Q48K, and S20N/Q48K, antiviral activity increased to around 13-fold which is still less than the node 1 ancestor (17-fold; Figure 5B). Similarly, patas monkey A3H with single mutations at sites 25 (R25P) and 51 (E51K) did not improve restriction, while a H17R mutation slightly increased the antiviral activity compared to wild-type (3-fold to 5-fold; Figure 5B). However, this single mutation at position 17 did not make patas monkey A3H comparable to the node 1 ancestor. Inserting both H17R and R25P mutations in patas monkey A3H similarly did not further improve restriction (Figure 5B). Taken together, the inability of single point mutations to restore antiviral activity to its ancestral state emphasizes that changes at multiple sites have functional consequences in AGM and patas monkey A3H.

Of interest, site 48 is the only residue that is fixed in all AGMs sequenced for this study. This demonstrates that it occurred first, whereas the other sites are polymorphic and have not yet become fixed in the species (Table 1). To determine whether site 48 alone is sufficient to decrease antiviral activity we added a K48Q mutation in the node 1 background and tested its antiviral activity. However, node 1 K48Q has similar antiviral activity and expression (Figure 5C, bottom), showing that epistasis between multiple amino acids may play an essential role for viral restriction by A3H.

Overall, these data establish that amino acids in the N-terminal portion of A3H are important for antiviral activity. While no tested combinations increase the antiviral activity of AGM or patas monkey A3H equivalent to the node 1 ancestor, all changes (minus one) are before the catalytic domain (Figure 5A). Two out of 3 residues responsible for the loss of activity in patas monkey A3H, sites 17 and 25, are also polymorphic in AGMs (Table 1), suggesting that the loss began at a shared common ancestor not sampled by our analysis. Significantly, many Old World monkeys encode additional mutations near amino acid 15, whose loss in human A3H is known to affect protein stability (13). For example, residues 18 – 23 have been deleted in De Brazza’s monkey (Figure 5A). Such a large deletion in this region would likely impact the ability of A3H to inhibit viral infection in this species. Similarly, talapoin A3H encodes a histidine at position 18 instead of an arginine, similar to AGM A3H, suggesting that its activity may also be lost (Figure 5A). In contrast, rhesus macaque and sooty mangabey A3H encode active ancestral amino acids at such residues, such as two arginines at positions 17 and 18 (Figure 5A), which may result in the potent antiviral activity of these modern proteins. This implies that loss of A3H function has been lost independently at various points of evolutionary history in both hominoids and Old World monkeys.

### Lack of evidence that evolution in A3H leading to AGM has been driven by Vif

The variability in A3H function throughout Old World monkey evolution suggests an outside selective pressure is driving its loss. Antiviral restriction factors are often rapidly evolving and undergo positive selection (38), which is defined by an excess of nonsynonymous mutations compared to synonymous mutations. Evolutionary conflicts between host restriction factors and viral proteins to either maintain or escape interactions result in an accumulation of nonsynonymous mutations at binding interfaces. A previous study found that A3H is under positive selection (14). However, the study used fewer primate sequences which can bias the analysis. Therefore, we wanted to re-test positive selection in primate A3H using additional sequences we obtained from Old World monkeys. Using the PAML (phylogenetic analysis by maximum likelihood) program (39), we calculated the number of nonsynonymous mutations (dN) over the number of synonymous mutations (dS) for the entire *A3H* gene as well as individual codons. In agreement with previous data, we found that the *A3H* gene is under positive selection and with a dN/dS ratio of 1.3 in all primates. Additionally, models that allow codons to evolve under positive selection fit the data significantly better than models that do not for all primate clades (Figure 6A). We evaluated individual sites within the gene and found that in all primates, a total of 19 residues are under positive selection with a posterior probability > 0.98. One site, position 90, has been implicated in Vif-binding interactions, but only had a posterior probability > 0.98 in one codon model (F3×4; Figure 6A). Since selection can sometimes be driven by the inclusion of specific clades in the analysis, we also condensed our analysis to Old World monkeys alone and found that 5 sites remain under positive selection (blue arrows, Figure 6B), all of which are polymorphic in AGMs (Table 1). No positively selected residues in Old World monkey A3H are in the putative Vif-binding region (35, 36). Thus, we conclude that multiple sites are under diversifying selection in primates but are likely not driven by lentiviral Vif.

**Figure 6.**
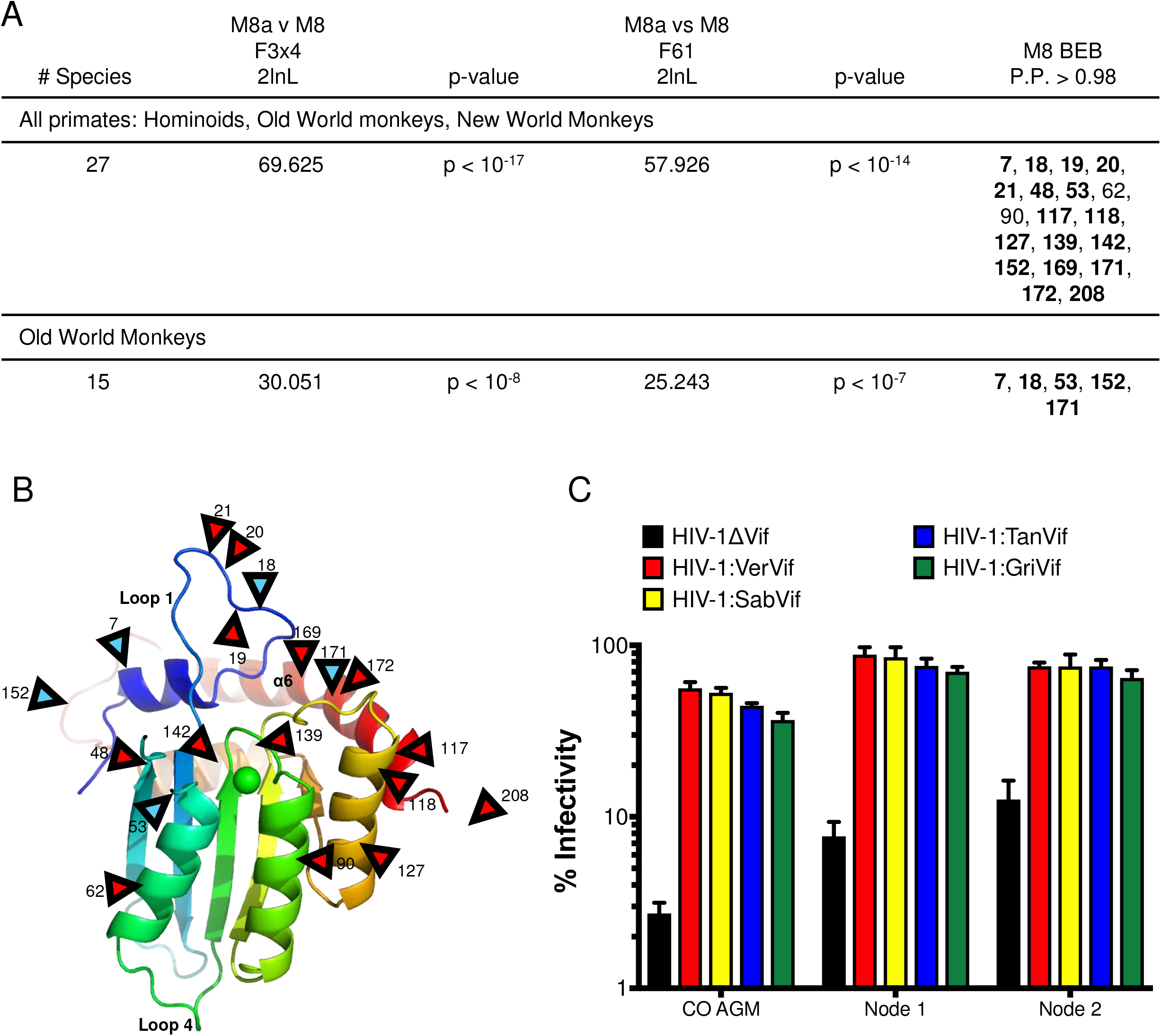
Evolution of A3H is not driven by Vif. **A**. Results of positive selection analyses of primate A3H. The last column lists sites under positive selection with dN/dS >1 with a posterior probability > 0.98 under M8 Bayes Empirical Bayes (BEB) implemented in PAML model 8. Sites are relative to African green monkey A3H. Sites that had a posterior probability > 0.98 in both codon models (F3×4 and F61) are bolded. **B**. A3H from individual PR01190 was modeled onto the pig-tailed macaque A3H structure previously described by Bohn et al, 2017 (PDB 5W3V). Locations of positively selected sites are denoted by red triangles. Blue triangles denote sites also under positive selection in Old World primates. Site 208 is part of a region deleted in the crystallized protein and is therefore not resolved in the model. **C**. Single-cycle infectivity assay done with *HIVΔvif* and HIV-1 expressing SIVagm Vif in the presence of codon-optimized AGM A3H haplotype 1 (CO AGM), node 1 ancestor, and node 2 ancestor. Relative infection was normalized to viral infectivity in the absence of A3 proteins. Averages of three replicates, each with triplicate infections ( ± SEM) are shown.

In order to test the hypothesis that the changes in A3H in the lineages leading to the AGMs were not driven by the Vif protein of the lentiviruses that infect AGMs, we tested the restriction capabilities of the node 1 and node 2 ancestors against HIV-1 proviruses expressing SIVagm Vif. We also included the codon-optimized AGM A3H since, as it is isolated from an AGM, should be susceptible to degradation by all Vifs encoded by SIVagm. Of interest, the node 2 ancestor encodes an aspartic acid (D) at amino acid 100, which has previously been important for differences in Vif-biding for SIVcpz and HIV-1, while the node 1 ancestor and all AGM A3Hs encode an asparagine (N) (35, 36). The antiviral activity of all ancestors, as well as codon-optimized AGM A3H, was counteracted by each Vif, resulting in a rescue of viral infection (Figure 6C). Because there are no differences in the ability of Vif to rescue viral infection, these data suggest that Vif is not the primary force on A3H evolution in AGMs, which is instead driven by different selective pressures.

## Discussion

We show evidence for the recurrent functional loss of APOBEC3H in primates. We found that a decrease in protein expression levels, as well as amino acid mutation in the N-terminal region, results in lower antiviral activity. Using molecular reconstruction of ancestral *A3H* sequences, we found that the most recent common ancestor of AGMs and patas monkeys likely encoded an active A3H, similar to other common ancestors throughout evolutionary history. This suggests that the recurrent loss is a more recent event in primate evolution. Selective pressure by Vif is does not appear to be a primary force behind the evolution of A3H in the AGM clade, but, as loss has occurred both in humans and in other Old world monkeys, there may be a fitness cost to encoding this mutator protein over long evolutionary time periods.

### Molecular evolution of A3H protein impacts expression levels and antiviral activity

While increasing the amount of A3H present in cells did increase its capability to inhibit viral infection, greater antiviral activity does not perfectly correlate with higher protein expression. Surprisingly, codon-optimization increased both the expression level and antiviral activity of A3H. However, codon-optimized AGM A3H haplotype 1 was not as potent as codon-optimized rhesus macaque A3H, demonstrating that both protein expression level and amino acid differences have functional consequences (Figure 3A and B). In support of this idea, while AGM A3H haplotypes 1 and 8 have the fewest amino acid changes from the most recent common ancestor with another species (Table 2, Figure 5A). Multiple mutations are required to allow the inactive extant protein to act like its active ancestor. Position 48 is fixed in AGMs, indicating that this change likely occurred first, while subsequent amino acid changes concurrently altered the antiviral activity of the protein. However, a K48Q mutation in the node 1 ancestor does not decrease its function (Figure 5C), demonstrating that epistasis between observed amino acid changes more likely lead to its loss. For instance, the AGM A3H double mutant H18R/Q48K inhibits viral infection more comparable to the node 1 ancestor while the individual mutants do not, indicating that both residues are required to gain antiviral activity (Figure 5B). Other AGM A3H haplotypes have also accumulated additional nonsynonymous mutations (Figure 2A), indicating that additional genetic drift may be actively driving A3H to become less antiviral in AGMs, as many animals encode proteins with different expression levels and antiviral activity (Figure 2B and C).

### Why has primate A3H maintained partial activity rather than a complete loss?

Loss of protein activity can be indicative of a fitness cost. In the case of human A3H, haplotypes encoding unstable proteins have been linked to greater cancer risk, likely due to its nuclear localization and proximity to host DNA (19). Moreover, human A3H haplotypes III and IV, which have a deletion at amino acid 15, have little to no antiviral activity. We observe that AGM A3H haplotypes vary greatly in their antiviral restriction and can restrict viral infection anywhere from 17 – 70% (Figure 2A).

However, despite this immense variation in AGM A3H antiviral activity, we observed no large deletions or premature stop codons, which would indicate that the gene itself is being lost. Why has the antiviral activity of A3H not been lost completely in AGMs? It is possible that A3H has been co-opted for a non-antiviral function in Old World monkeys, thus it is preserved. Alternatively, A3H could be retained through linkage to protective A3G haplotypes. A3G and A3H are located close together on chromosome 19 in AGMs, meaning A3H haplotypes that have ultimately lost most of their function may still be passed onto offspring, particularly if the A3G allele encodes a protein providing a selective advantage. In support of this idea, we noticed poorly active A3H proteins in a monkey encoding protective A3G, as characterized in a separate study (8). This individual, V005, encodes A3H haplotype 11 as well as an A3G haplotype that cannot be antagonized by Vif proteins from SIVagm.ver or SIVagm.tan. Since A3G is a more potent antiviral and this protein protects individuals from two strains of SIVagm, the protection it supplies may supersede any detrimental effects incurred by the presence of an inactive A3H haplotype.

### Loss of activity of A3H due to recurrent mutations in a putative RNA binding domain

Two mutations in human A3H gave rise to inactive protein variants: R105G and a deletion of amino acid 15. Of interest, amino acid 15 is positioned within loop 1 of the A3H protein. Recent work has revealed that loop 1, 7, and α6 are important for binding to an RNA duplex (32–34). We similarly identified that amino acids 18 and 20, found within loop 1, are important for increasing the antiviral activity of AGM A3H (Figure 5B). The analogous locations of such residues may suggest that these changes impact the antiviral activity of A3H by affecting its capability to bind to viral RNA. Dual amino acid changes at residues 18 and 20 in AGMs increase the antiviral activity of A3H close to the levels of a recent common ancestor. An additional mutation close to the catalytic site, Q48K, together with either H18R or S20N in AGMs, further increased antiviral activity. Furthermore, we found that amino acid 17 similarly increased the antiviral activity of patas monkey A3H. This residue is polymorphic in AGMs, suggesting that this change may have occurred in a common ancestor not tested in this study. We also find that other Old World monkeys have changes within loop 1, such as a six amino acid deletion of residues 18 – 23 in De Brazza’s monkey and an R18H mutation in talapoin (Figure 5A). The diversity within loop 1 of Old World monkey A3H is indicative that A3H activity was possibly lost multiple times independently. RNA-binding has been shown to play an important role in the antiviral activity of A3H (32, 33) and thus loss of RNA-binding may result in functional loss.

### Vif does not appear to play a role in evolution of A3H along lineage leading to AGMs

AGMs are highly polymorphic in a number of antiviral genes that are specific to infection with lentiviruses in both AGMs explicitly and other primates (31), further highlighting the adaptation to SIV within this particular species. Indeed, we found that multiple sites in A3H are under positive selection in primates, which is suggestive of an evolutionary arms race between A3H and another protein. Evidence of A3H-Vif interactions are evident in hominoids, as SIVcpz Vif adaptation to stable human A3H was crucial for transmission of SIVcpz (11) to humans and HIV-1 Vif is highly variable in its ability to target stable human A3H proteins (17, 21–23). Interactions between Vif and other A3 proteins have also been well-characterized in chimpanzees and humans (7, 35, 36). Multiple A3s in chimpanzees protect them from infection by SIVs in Old World monkeys. Therefore, Vif must adapt to antagonize multiple A3 proteins such as A3D and A3G (10). However, we find that Vif did not play a role in A3H’s evolution in AGMs since ancestral proteins were comparably susceptible to antagonism by Vif. Loss of A3H function may facilitate cross-species transmission of SIVagm strains between AGM subspecies (40) due a diminished A3 repertoire. Although many A3 proteins in the family may be redundant, encoding a diverse range of A3s is likely important to achieve maximum protection against lentiviruses.

Since lentiviruses have infected simian primates for millions of years (7), it is unlikely that the lack of residues altering A3H-Vif interactions in AGMs stems from recent infection of Old World monkeys. It is possible that the changes are the result of genetic drift or driven by a different viral pathogen in these primates. Conversely, positively selected residues may indicate evolutionary toggling to preserve or eliminate protein function. This is supported by the finding that many amino acid residues found to increase the antiviral activity of A3H in AGMs are under positive selection (Figure 6A, 6B). It is also possible that the relative importance of different A3 proteins may change dependent on the evolutionary history of a species, driven by the redundancy of the protein family. In the Old World monkeys studied here, inactive A3H proteins may impart an increased risk for host genome mutations, thus its function was lost due to the balance between viral protection and host fitness.

Our data implies that A3H function was lost prior to the divergence of different SIVagm strains due to its inactivity in all AGMs, not just specific subspecies. If this is the case, Vif proteins from the ancestral virus may not have required adaptation to escape the antiviral activity of A3H and instead evolved in response to pressure from the more potent A3G. Expansion of the primate *A3* locus provides flexibility of this antiviral protein family to take different trajectories throughout evolution. We have demonstrated that A3H activity is fluid throughout the evolutionary history of primates. In addition to previous work in humans, the A3H homolog in felines, APOBEC3Z3, was recently shown have a similar functional loss (41), demonstrating that loss of A3H and its homologs are frequent throughout a variety of animal species. Such widespread loss of function is suggestive of a potential fitness cost to hosts, although the presence of modern and active A3H proteins exemplifies the importance of encoding a diverse *A3* locus in primates.

## Materials and methods

### APOBEC3H cDNA Amplification and Sequencing

APOBEC3H cDNAs were cloned by nested RT-PCR (QIAGEN One-step RT PCR Kit) or PCR (Accuprime Pfx) from RNA or gDNA isolated from AGM peripheral blood mononuclear cells or cell lines. Sample origins and extractions have been previously described (8, 30). For each sample, PCR products were amplified and sequenced using primers designed to amplify African green monkey A3H (Forward: CACGAATTCGCCACCATGTATCCATACGATGTTCCAGATTACGC TGCTCTGCTAACAGCCAAA Reverse: CACGAGCTCATCTTGAGTTGAGTGT). Primers for gDNA amplification were designed to target intronic regions. Heterozygous sequences were cloned using pGEM T-Easy vector system (Promega) and TOPO TA cloning kit (Invitrogen) to analyze individual clones. Additional primate sequences were obtained from gDNA isolated from immortalized cell lines using QIAGEN DNeasy Blood & Tissue Kit. The following cell lines from Coriell Cell Repositories (Camden, NJ) were used: patas monkey *(Erythrocebus patas;* ID no. 6254), De Brazza’s monkey *(Cercopithecus neglectus;* PR01144), Wolf’s guenon *(Cercopithecus wolfi;* PR01241), mustached guenon *(Cercopithecus cephus,* PR00527), Allen’s swamp monkey *(Allenopithecus nigroviridis;* PR00198), and Francois’ Leaf monkey *(Trachypithecus francoisi;* PR01099). Talapoin *(Miopithecus talapoin;* OR755) cells were obtained from Frozen Zoo (San Diego, CA).

### Expression Constructs and Plasmids

Primate *A3H* genes were cloned from cDNA. A 5’ hemagluttinin (HA) tag was added via PCR (Forward:

GTGGTGGAATTCATGTATCCATACGATGTTCCAGATTACGCTGCTCTGCT Reverse: CTAGACTCGAGTCATCTTGAGTT). The products were digested using EcoRI/XhoI restriction enzymes and ligated into a mammalian pcDNA 3.1 vector (Invitrogen, #V79020). Site-directed mutagenesis was completed with the QuikChange II Site-Directed Mutagenesis Kit (Agilent, #200524) to construct all ancestral genes and mutants. The *A3H* gene from patas monkeys was generated by gene synthesis (IDT) and cloned into the pcDNA 3.1 backbone. *A3H* genes from AGM and rhesus macaque were codon-optimized based on usage frequencies in primates (human and rhesus ‘ macaque) to remove rare codons within the gene in Geneious (Biomatters Ltd.). Codon-optimized sequences were generated by gene synthesis and cloned into pcDNA 3.1.

### Cell Lines, Transfections, and Western Blot Analysis

HEK293T, HeLa, and Cos7 cell lines (ATCC) were maintained in Dulbecco’s modified Eagle’s medium (DMEM) with 10% fetal bovine serum (Corning, #35-015-CV) and 100 μg/mL penicillin/streptomycin (Gibco, #15140-122) at 37°C. SupT1 cells (ATCC) were maintained in RPMI 1640 with 10% fetal bovine serum and 100 μg/mL penicillin/streptomycin in the same conditions. Transfections were done in serum-free DMEM with TransIT-LT1 transfection reagent (Mirus Bio, #MIR 2305) at a ‘ reagent:plasmid DNA ratio of 3:1. For western blot analysis, cells were lysed in ice-cold NP40 buffer (0.5% NP40, 20mM NaCl, 50mM Tris pH 7.5) with protease inhibitors (Roche Complete Mini, EDTA-free tablets, #11836170001). Lysates were quantified using a Pierce BCA Protein Assay Kit (Thermo Scientific, #23225) and 10 μg of protein was resolved by SDS-PAGE, transferred to a PVDF membrane, and probed with anti-HA (BioLegend, #901503) and anti-actin (Sigma, #A2066) or anti-vinculin (Proteintech, #66305-1) antibodies at a 1:2000 dilution. Anti-mouse or anti-rabbit secondary antibodies were used at a 1:5000 dilution (Santa Cruz Biotechnology, sc-2005, sc-2004).

### Immunofluorescence

HeLa cells were seeded onto 18mm (VWR, #48380 046) coverslips seeded with 4 × 10^4^ cells/mL and transfected with 500 ng of A3H-expressing plasmids the next day. 48 hours after transfection, the coverslips were fixed in 2% paraformaldehyde, permeabilized in 0.5% PBS/Triton-X, and blocked in PBS/BGS. HA-tagged proteins were detected using the same HA antibody used for western blots at a 1:1000 dilution followed by an anti-mouse AF488 antibody at 1:400 (Invitrogen, #A11001). Nuclei were stained in SlowFade Gold antifade reagent with DAPI mounting media (Life Technologies, #S36939). Images were taken on a Nikon E800 microscope.

### Phylogenetic Analysis

*A3H* AGM sequences were analyzed phylogenetically using a Bayesian Monte Carlo Markov chain (MCMC) approach implemented in BEAST v1.7.1. Sequence alignments were constructed using MAFFT align function in Geneious (Biomatters Ltd.) and underwent 10,000,000 MCMC generations using HKY85 substitution model, gamma site heterogeneity model, estimated base frequencies, and constant population size coalescent as the tree prior.

### Positive Selection Analysis

The 27 primate *A3H* sequence alignment was analyzed using HyPhy GARD analysis to ensure there was no recombination in the gene (42). The species phylogeny (27) was input into the CODEML sites model of PAML (39) along with the nucleotide alignment to detect positive selection at individual sites. The p-value was calculated by twice the difference in log-likelihood between models M7 and M7 as well as M8 and M8a with two degrees of freedom. Analysis was conducted with both the F3×4 and F61 codon frequency models with omega values of 0.4 and 1.5. Data from F3×4 and F61 models are shown in Table 1. Positively selected sites were categorized as those with an M8 Bayes Empirical Bayes posterior probability greater than 98%.

### Ancestral Reconstruction

The ancestors at specific nodes within the Old World monkey clade were reconstructed using the FASTML webserver (fastml.tau.ac.il; last accessed March 2018; (37)). The 27 primate *A3H* sequence alignment was used in conjunction with the species tree to generate marginal reconstructions of codon sequences.

### Single-Cycle Infectivity Assays

HEK293T cells were plated in 1mL in 12-well plates at 1.25 × 10^5^ cells/mL. After cells reached between 50 – 70% confluency, they were co-transfected with 250 ng of A3H or empty expression plasmid, 600 ng of proviral plasmid, and 100 ng of L-VSV-G (vesicular stomatitis virus glycoprotein) for pseudotyping in 100 μL serum-free medium with TransIT-LT1 transfection reagent (Mirus Bio). Supernatants containing virus were harvested after 48 hours and clarified through 0.2-micron filters. Viral titers were determined by measuring reverse-transcriptase (RT) activity by qPCR as described previously (43). In short, viral supernatants were lysed in 2X lysis buffer (0.25% Triton X-100, 50 mM KCl, 100 mM Tris-HCl, 40% glycerol) in the presence of 4U RNase inhibitor (Fermentas, #EO0382). qRT-PCR reactions were set up with an MS2 RNA template using the Takyon Rox SYBR MasterMix dTTP Blue kit (Eurogentec, #UF-RSMT-B0101) alongside a standard curve made with a stock virus of previously determined titers. The primers used to amplify duplicate reactions were: TCCTGCTCAACTTCCTGTCGAG (forward) and CACAGGTCAAACCTCCTAGGAATG (reverse). qRT-PCR was performed on an ABI QuantStuido5 Real Time PCR machine. 2000 mU/mL was used to infect SupT1 cells plated at 2 × 10^4^ cells/well in a 96-well plate in media supplemented with 20 μg/mL DEAE-Dextran. Infections were done in triplicate for 48 – 64 hours. Luciferase activity was measured with Bright-Glo Luciferase Assay Reagent (Promega) on a LUMIstar Omega luminometer.

### Accession Numbers

The GenBank accession numbers for new Old World monkey *A3H* sequences, including De Brazza’s, Wolf’s Guenon, mustached guenon, patas monkey, talapoin, Francois’ Leaf monkey, Allen’s swamp monkey, and AGM haplotype, reported here are MH231602 – MH231609.

## Acknowledgements

We thank the following investigators for providing samples used in this study: M. Muller-Trutwin (AGM PBMC), C. Apetrei (AGM PBMC and SIVagm.Sab92018), and N. Freimer and A. Jasinska (AGM DNA). We additionally thank the National Institutes of Health (NIH) Nonhuman Primate Research Resource for the V038 and AG23 AGM cells as well as SIVagm.Gri+ plasma. African sabaeus samples used in this study were collected as a part of the Systems Biology Sample Repository funded by NIH Grants R01RR016300 and R0100010980, with the particular sabaeus samples used in this study through the Department of Parks and Management Ministry of Forestry and the Environment and Medical Research Council Unit, The Gambia. We thank the Fred Hutchinson Cancer Research Center Shared Resources Genomics Core for sequencing. We thank Ferdinand Roesch, Nicholas Chesarino, Mollie McDonnell and Emily Hsieh for constructive comments on the manuscript. We also thank Harmit Malik, Rick McLaughlin, and Rossana Colon-Thillet for helpful discussions. This work was supported by NIH grant R01 AI30927 (M.E) and the Viral Pathogenesis Training Grant T32AI083203 (E.G).

## References

1. Harris RS, Dudley JP. 2015. APOBECs and virus restriction. Virology 479–480:131–145.

2. Harris RS, Liddament MT. 2004. Retroviral restriction by APOBEC proteins. Nat Rev Immunol 4:868–877.

3. Stavrou S, Ross SR. 2015. APOBEC3 Proteins in Viral Immunity. J Immunol 195:4565–4570.

4. Hultquist JF, Lengyel JA, Refsland EW, LaRue RS, Lackey L, Brown WL, Harris RS. 2011. Human and rhesus APOBEC3D, APOBEC3F, APOBEC3G, and APOBEC3H demonstrate a conserved capacity to restrict Vif-deficient HIV-1. J Virol 85:11220–11234.

5. Belanger K, Savoie M, Rosales Gerpe MC, Couture JF, Langlois MA. 2013. Binding of RNA by APOBEC3G controls deamination-independent restriction of retroviruses. Nucleic Acids Res 41:7438–7452.

6. Simon V, Bloch N, Landau NR. 2015. Intrinsic host restrictions to HIV-1 and mechanisms of viral escape. Nat Immunol 16:546–553.

7. Compton AA, Emerman M. 2013. Convergence and divergence in the evolution of the APOBEC3G-Vif interaction reveal ancient origins of simian immunodeficiency viruses. PLoS Pathog 9:e1003135.

8. Compton AA, Hirsch VM, Emerman M. 2012. The host restriction factor APOBEC3G and retroviral Vif protein coevolve due to ongoing genetic conflict. Cell Host Microbe 11:91–98.

9. McCarthy KR, Kirmaier A, Autissier P, Johnson WE. 2015. Evolutionary and Functional Analysis of Old World Primate TRIM5 Reveals the Ancient Emergence of Primate Lentiviruses and Convergent Evolution Targeting a Conserved Capsid Interface. PLoS Pathog 11:e1005085.

10. Etienne L, Bibollet-Ruche F, Sudmant PH, Wu LI, Hahn BH, Emerman M. 2015. The Role of the Antiviral APOBEC3 Gene Family in Protecting Chimpanzees against Lentiviruses from Monkeys. PLoS Pathog 11:e1005149.

11. Zhang Z, Gu Q, de Manuel Montero M, Bravo IG, Marques-Bonet T, Haussinger D, Munk C. 2017. Stably expressed APOBEC3H forms a barrier for cross-species transmission of simian immunodeficiency virus of chimpanzee to humans. PLoS Pathog 13:e1006746.

12. Bishop KN, Holmes RK, Sheehy AM, Davidson NO, Cho SJ, Malim MH. 2004. Cytidine deamination of retroviral DNA by diverse APOBEC proteins. Curr Biol 14:1392–1396.

13. OhAinle M, Kerns JA, Li MM, Malik HS, Emerman M. 2008. Antiretroelement activity of APOBEC3H was lost twice in recent human evolution. Cell Host Microbe 4:249–259.

14. OhAinle M, Kerns JA, Malik HS, Emerman M. 2006. Adaptive evolution and antiviral activity of the conserved mammalian cytidine deaminase APOBEC3H. J Virol 80:3853–3862.

15. Wang X, Abudu A, Son S, Dang Y, Venta PJ, Zheng YH. 2011. Analysis of human APOBEC3H haplotypes and anti-human immunodeficiency virus type 1 activity. J Virol 85:3142–3152.

16. Harari A, Ooms M, Mulder LC, Simon V. 2009. Polymorphisms and splice variants influence the antiretroviral activity of human APOBEC3H. J Virol 83:295–303.

17. Refsland EW, Hultquist JF, Luengas EM, Ikeda T, Shaban NM, Law EK, Brown WL, Reilly C, Emerman M, Harris RS. 2014. Natural polymorphisms in human APOBEC3H and HIV-1 Vif combine in primary T lymphocytes to affect viral G-to-A mutation levels and infectivity. PLoS Genet 10:e1004761.

18. Li MM, Emerman M. 2011. Polymorphism in human APOBEC3H affects a phenotype dominant for subcellular localization and antiviral activity. J Virol 85:8197–8207.

19. Starrett GJ, Luengas EM, McCann JL, Ebrahimi D, Temiz NA, Love RP, Feng Y, Adolph MB, Chelico L, Law EK, Carpenter MA, Harris RS. 2016. The DNA cytosine deaminase APOBEC3H haplotype I likely contributes to breast and lung cancer mutagenesis. Nat Commun 7:12918.

20. Burns MB, Temiz NA, Harris RS. 2013. Evidence for APOBEC3B mutagenesis in multiple human cancers. Nat Genet 45:977–983.

21. Li MM, Wu LI, Emerman M. 2010. The range of human APOBEC3H sensitivity to lentiviral Vif proteins. J Virol 84:88–95.

22. Ooms M, Brayton B, Letko M, Maio SM, Pilcher CD, Hecht FM, Barbour JD, Simon V. 2013. HIV-1 Vif adaptation to human APOBEC3H haplotypes. Cell Host Microbe 14:411–421.

23. Ooms M, Letko M, Binka M, Simon V. 2013. The resistance of human APOBEC3H to HIV-1 NL4–3 molecular clone is determined by a single amino acid in Vif. PLoS One 8:e57744.

24. Naruse TK, Sakurai D, Ohtani H, Sharma G, Sharma SK, Vajpayee M, Mehra NK, Kaur G, Kimura A. 2016. APOBEC3H polymorphisms and susceptibility to HIV-1 infection in an Indian population. J Hum Genet 61:263–265.

25. Sakurai D, Iwatani Y, Ohtani H, Naruse TK, Terunuma H, Sugiura W, Kimura A. 2015. APOBEC3H polymorphisms associated with the susceptibility to HIV-1 infection and AIDS progression in Japanese. Immunogenetics 67:253–257.

26. Fregoso OI, Ahn J, Wang C, Mehrens J, Skowronski J, Emerman M. 2013. Evolutionary toggling of Vpx/Vpr specificity results in divergent recognition of the restriction factor SAMHD1. PLoS Pathog 9:e1003496.

27. Perelman P, Johnson WE, Roos C, Seuanez HN, Horvath JE, Moreira MA, Kessing B, Pontius J, Roelke M, Rumpler Y, Schneider MP, Silva A, O’Brien SJ, Pecon-Slattery J. 2011. A molecular phylogeny of living primates. PLoS Genet 7:e1001342.

28. Wertheim JO, Worobey M. 2007. A challenge to the ancient origin of SIVagm based on African green monkey mitochondrial genomes. PLoS Pathog 3:e95.

29. Peeters M, Courgnaud V. 2002. Overview of Primate Lentiviruses and Their Evolution in Non-human Primates in Africa. HIV Sequence Compendium 2002:2–23.

30. Spragg CJ, Emerman M. 2013. Antagonism of SAMHD1 is actively maintained in natural infections of simian immunodeficiency virus. Proc Natl Acad Sci U S A 110:21136–21141.

31. Svardal H, Jasinska AJ, Apetrei C, Coppola G, Huang Y, Schmitt CA, Jacquelin B, Ramensky V, Muller-Trutwin M, Antonio M, Weinstock G, Grobler JP, Dewar K, Wilson RK, Turner TR, Warren WC, Freimer NB, Nordborg M. 2017. Ancient hybridization and strong adaptation to viruses across African vervet monkey populations. Nat Genet 49:1705–1713.

32. Ito F, Yang H, Xiao X, Li SX, Wolfe A, Zirkle B, Arutiunian V, Chen XS. 2018. Understanding the Structure, Multimerization, Subcellular Localization and mC Selectivity of a Genomic Mutator and Anti-HIV Factor APOBEC3H. Sci Rep 8:3763.

33. Shaban NM, Shi K, Lauer KV, Carpenter MA, Richards CM, Salamango D, Wang J, Lopresti MW, Banerjee S, Levin-Klein R, Brown WL, Aihara H, Harris RS. 2018. The Antiviral and Cancer Genomic DNA Deaminase APOBEC3H Is Regulated by an RNA-Mediated Dimerization Mechanism. Mol Cell 69:75–86 e79.

34. Bohn JA, Thummar K, York A, Raymond A, Brown WC, Bieniasz PD, Hatziioannou T, Smith JL. 2017. APOBEC3H structure reveals an unusual mechanism of interaction with duplex RNA. Nat Commun 8:1021.

35. Nakashima M, Tsuzuki S, Awazu H, Hamano A, Okada A, Ode H, Maejima M, Hachiya A, Yokomaku Y, Watanabe N, Akari H, Iwatani Y. 2017. Mapping Region of Human Restriction Factor APOBEC3H Critical for Interaction with HIV-1 Vif. J Mol Biol 429:1262–1276.

36. Ooms M, Letko M, Simon V. 2017. The Structural Interface between HIV-1 Vif and Human APOBEC3H. J Virol 91.

37. Ashkenazy H, Penn O, Doron-Faigenboim A, Cohen O, Cannarozzi G, Zomer O, Pupko T. 2012. FastML: a web server for probabilistic reconstruction of ancestral sequences. Nucleic Acids Res 40:W580–584.

38. Daugherty MD, Malik HS. 2012. Rules of engagement: molecular insights from host-virus arms races. Annu Rev Genet 46:677–700.

39. Yang Z. 2007. PAML 4: phylogenetic analysis by maximum likelihood. Mol Biol Evol 24:1586–1591.

40. Bell SM, Bedford T. 2017. Modern-day SIV viral diversity generated by extensive recombination and cross-species transmission. PLoS Pathog 13:e1006466.

41. Konno Y, Nagaoka S, Kimura I, Takahashi Ueda M, Kumata R, Ito J, Nakagawa S, Kobayashi T, Koyanagi Y, Sato K. 2018. A naturally occurring feline APOBEC3 variant that loses anti-lentiviral activity by lacking two amino acid residues. J Gen Virol doi:10.1099/jgv.0.001046.

42. Kosakovsky Pond SL, Posada D, Gravenor MB, Woelk CH, Frost SD. 2006. GARD: a genetic algorithm for recombination detection. Bioinformatics 22:3096–3098.

43. Vermeire J, Naessens E, Vanderstraeten H, Landi A, Iannucci V, Van Nuffel A, Taghon T, Pizzato M, Verhasselt B. 2012. Quantification of reverse transcriptase activity by real-time PCR as a fast and accurate method for titration of HIV, lenti- and retroviral vectors. PLoS One 7:e50859.

